# Differential effects of Doxorubicin and Actinomycin D on the stability of RNA binding proteins, RBM10 and RBM5: Actinomycin D promotes the nuclear speckles targeting of RBM10 and RBM5 through the novel structural elements

**DOI:** 10.1101/627927

**Authors:** Koji Nishio, Shanlou Qiao, Kung Sang Chang

## Abstract

**Background:** RNA binding motif (RBM) proteins, RBM10v1, RBM10v2 and RBM5 share a high degree of the conserved domains. So far, the drug-sensitivities of the RBMs in tumor cells have not been fully examined.

**Objective:** The expression profiles of RBM10 and RBM5 in several virus-transformed tumor cells, and the effect of the most established antitumor agents, actinomycin D and doxorubicin, were investigated.

**Methods and Results:** Doxorubicin and actinomycin D differentially reduced RBM10 and RBM5 protein, respectively in both of HeLa and COS-7 cells. RBM10 protein was highly sensitive to doxorubicin in HeLa, COS-7 and A549 cells. In silico analysis revealed the several sumoylation sites of RBM10 and its sumoylated form could be targeted for the activated ubiquitin proteasome system. Actinomycin D affected the nuclear speckles localization of RBM10 and RBM5 in COS-7 and A549 lung carcinoma cells. Addition of actinomycin D in the culture medium and following culture for 3〜4 hours promoted the prominent nuclear speckles of RBM10v2-GFP and RBM5. Hence, we explored the subnuclear localization of the full length RBM10v2 (852aa) and the amino terminally truncated forms and the responsible structural elements. The amino terminally truncated RBM10v2 [#486-852, #642-852], RBM10v2 [#648-852], RBM10v2 [#681-759, #681-852], and RBM10v2 [#660-852] retained the targeting elements for the nuclear speckles, nucleoplasm, nucleoli and whole nuclei, respectively.

**Conclusion:** RBM10 is highly sensitive to doxorubicin. Actinomycin D affects the structural elements of RBM10 and promotes the nuclear speckles targeting. The C-terminal regions: RBM10v2 [#642-647], [#642-659], and [#660-680] play critical roles in the targeting to the subnuclear compartments. SIM and sumoylation sits of RBM10 and PML4 are important for molecular interaction of RBM10 and PML.

## Introduction

Nuclear speckles enrich in the pre-mRNA splicing regulators and organize active genes (1). The initial step of splicing is binding of U1 snRNP with the pre-mRNA to form spliceosomal E complex (2). RBM5 and RBM10 assemble with the stalled spliceosome A complex and precatalytic B complexes, respectively (3). The precatalytic spliceosome B complex assembles with preformed U4/U6/U5 tri-snRNP. Importantly, ubiquitylation and sumoylation involve in the modification and assembly of spliceosomal components (4,5).

RBM10 is proposed as an important regulator of alternative splicing (6). The human RBM10 gene is located at a region of chromosome X, p11.3-p11.23 and frequently mutated in lung adenocarcinomas (7). Frame-shift mutations of the RBM10 gene cause X-linked disorders in affected males (8,9). The human RBM5 gene is located at a region of chromosome 3, 3p21.3 which is frequently deleted in a variety of cancers (10). RBM5 or RBM10 involves in alternative splicing of Fas pre-mRNA (11,12). RBM10 pre-mRNA is alternatively spliced to RBM10v1 (930 AA) and RBM10v2 (852AA). The two RBM10 variants and RBM5 (815AA) share significant sequence similarities (13).

Doxorubicin (DOX), a toposiomerase II inhibitor inhibits the synthesis of DNA and RNA, activates caspase-2 and caspase-3 (14–17), and triggers the activation of ubiquitin proteasome system (18). Actinomycin D (AcD) inhibits the processes of RNA elongation and DNA replication, and activates several caspases (19–21). DOX generates free radicals and damages cardiac cellular membranes (22). In addition, DOX and a low concentration of AcD are known as nucleolar stress inducers (23). AcD promotes the nuclear speckles-targeting of RBM10, RBM6, SF3b^155^ and SR related protein (24–27). So far, the effects of doxorubicin and AcD on the expression of the RBMs have not been fully investigated.

HeLa and COS-7cells are hyperproliferative and known as SV40- and HPV-18-transformed cells, respectively. In the present study, protein levels of RBM10 and RBM5 were examined in HeLa, COS-7, A549 lung carcinoma cells, differentiated HK-2 cells or human normal fibroblasts TIG3S. These cells were subdivided into three groups according to the RBM protein levels. Interestingly, HeLa and COS-7 cells were regarded as high RBM-expressing, and A549 and TIG3S cells were regarded as low RBM-expressing, whereas HK-2 cells displayed the lowest expression profiles.

The antitumor agent sensitivities of RBM proteins were explored in HeLa, COS-7 and A549 cells. At first, the reductions of RBM10 and RBM5 proteins were differentially triggered by DOX and AcD, respectively. The proteolytic sensitivity of RBM10 following from the activation of ubiquitin-proteasome system will be discussed. AcD significantly promoted the targeting of RBM10 and RBM5 to the enlarged nuclear speckles. Finally, the novel structural elements which target RBM10 to the nuclear speckles, nucleolar or PML-nuclear bodies are proposed.

## Materials and Methods

### Cell lines and cell culture

COS-7 and HeLa PS (3) cells were provided from JCRB (Japanease Cancer Research Resources Bank, Tokyo, Japan). COS-7 cells represent SV40-large T antigen transformed immortal cell line. A549 human lung cancer cells were purchased from ATTC. HK-2, a cell line of human kidney proximal tubule was provided by Dr. Iwayama (Nagoya University, Nagoya, Aichi, Japan). HK-2 cells proliferate EGF-dependently and remain a functional phenotype of proximal tubule cells (28). HK-2 cells grew most slowly among the cells used. Fetal skin fibroblasts TIG3S at a 20 population doubling levels were gifted by Dr. Kondo (Tokyo Metropolitan Institute of Gerontology, Tokyo, Japan). These cells were cultured in 6-well plates, in which each well contained 2 ml of Dolbeco’s modified essential medium (DMEM) supplemented with 10% FCS, 1 mM glutamine and 60 μg/ml kanamycine and passaged at a split ratio of 1:2 or 1:4. The cells were grown to nearly full-confluence and the whole cell lysates (10 µg in SDS sample buffer) were prepared for Western blotting analysis of RBM10 and RBM5 protein. Total RNA was prepared with QIAGEN RNeasy mini kit and used for RT-PCR.

### Antibodies and Western blot analysis

Rabbit polyclonal anti-RBM10 against the N-terminal region (#1-166) of rat RBM10v2 was provided from Dr. Inoue (Osaka City University, Japan). The anti-RBM10 antibody specifically recognized RBM10v1 (130 kDa) and RBM10v2 (114kDa) (25). Rabbit polyclonal anti-RBM5 antibody against the C-terminal half (#408–815) of human RBM5 was provided from Dr. Oh (UCLA) (29). The cell lysates were separated by SDS 8%-polyacrylamide gel electrophoresis. The RBM proteins were examined with the primary antibodies and the following secondary anti-rabbit IgG conjugated with Horse radish peroxidase (A6154; Sigma-Aldrich) or conjugated with alkaline phasphatase (T2180;Applied Biosystems). β-actin was detected with mouse monoclonal anti-actin (A4700: Sigma-Aldrich) and anti-mouse IgG conjugated with alkaline phosphatase (A3688; Sigma-Aldrich). The specific protein bands were captured by Olympus μ-7000 digital camera and quantified by the image analysis software, Image-Pro plus (Media Cybernetics, MD. USA).

### Construction of expression vectors

An expression vector of human PML-Ⅳ(isoform 6, protein NP_002666) was provided by Dr Kung Sang Chang (University of Texas, USA). A GFP-fused construct of PML-Ⅵ (isoform1, GenBank at NCBI, accession no.AAA60125) was provided by Dr. Katano (National Institute of infectious diseases, Tokyo, Japan). Expression vectors of wild type human RBM5-GFP and human wild type RBM10v1 (930 amino acids) were provided by Dr. Valcárcel (University Pompeu Fabra, Spain) and by Kazusa DNA Research Center (Kisarazu, Chiba, Japan), respectively. Expression vectors of wild type rat RBM10v2 cDNA (852 amino acids), human RBM10v1-GFP and rat RBM10v2-GFP (25) were provided by Dr. Inoue (Osaka City University). RBM10 cDNA was amplified by polymerase chain reaction with high fidelity KOD DNA polymerase (Toyobo Inc., Japan). The amplified RBM10v1 and RBM10v2 were digested with Bgl II and EcoR I and then cloned into Bgl II and EcoR I sites of the RFP expression vector, pDsRed1-N1 (Clontech. Inc., USA). The C-terminal region of RBM10v2 [#486-852], [#642-852], [#648-852], [#660-852], [#648-759], [#681-852] or [#681-759] and the N-terminal region of RBM10v2 [#1-629] were amplified as described above and cloned into the pDsRed1-N1 and pNLS-EGFP-N3 vector, respectively. The coding sequences of RBM10 were confirmed by restriction digestion. The full length RBM10-GFP and RBM5-GFP proteins were detected with the specific antibodies.

### Antitumor agents, transfection and fluorescence microscopy

COS-7 or HeLa cells grown in 6-well plates were cultured in the presence or absence of doxorubicin (DOX) (44583: Sigma-Aldrich) or Actinomycin D (AcD) (A1410: Sigma-Aldrich) for 6 or 16 hours and the cell lysates were prepared.

A549 or COS-7 cells were cultured in Falcon 4-well culture slides (Lifetecnologies Inc. USA) until 80%-confulency was reached. Subconfluently grown A549 cells were exposed to DOX (0.5 µM or 1 µM) for 3 or 20 hours. The cells were fixed in 4% formalin-PBS for 10 minutes, washed in PBS and permealized in 1% Triton X-100-PBS for 5 minutes and stained with the primary anti-RBM5 or anti-RBM10 antibody and the Alexa^488^- or Alexa^555^-labeled anti-rabbit IgG.

COS-7 cells grown in an each well were transfected with 1μg of the expression vector of RBM10v2-RFP and 5μl of Lipofectoamin 2000 reagent, cultured for 48 hours at 37 ºC and exposed to 0.5 µM or 1 µM DOX for 8 hours. COS-7 cells were transfected with 1μg of the expression vector of RBM10v2-GFP as described above and exposed to 1 µM AcD for 4 or 20 hours. The nuclear speckles targeting of the RBMs were examined by Keyence BZ8000 fluorescence-phase contrast microscopy (Keyence Japan, Osaka).

COS-7 and A549 cells were transfected with 1μg of the expression vector of the N-terminally truncated RBM10v2-RFP, C-terminally truncated RBM10v2-GFP or RBM5-GFP, and exposed to 1 µM AcD for 3 hours, washed in PBS and fixed. Subnuclear compartments-targeting of the GFP (or RFP)-fused RBMs was examined by fluorescence-phase contrast microscopy as described above.

### Reverse transcription and PCR analysis

Reverse transcription was performed from 1 µg of total RNA using Expand reverse transcriptase (Roche) and oligo dT or random hexamer oligonucleotides as a primer. PCR was performed with GeneAmp PCR System 9700 (Applied Biosystems) using FastStart Taq DNA polymerase (Roche) as described in the manufacturer’s instruction. The PCR-reaction mixture contained 1x PCR buffer, with or without 1x GC-RICH solution supplemented with 2 mM MgCl_2_, 0.2 mM of each dNTP, 0.5 µM of each forward and reverse primers and 2 Units of FastStart Taq DNA polymerase. The cDNA was denatured at 95℃ for 6min and then amplified for 35 cycles using the following parameters: 95℃ for 30 sec, 55 ℃ for 30 sec and 1min for 72℃, and at 72 ℃ for 7 min (final extension). The endogenous mRNAs of human RBM10v1, RBM10v2, RBM5, Fas 5-6-7, Fas 5-7 and β-actin were amplified. To monitor the efficiency and reproducibility of PCR amplification, glycerolaldehyde 3-phosphate dehydrogenase (GAPDH) or β-actin cDNA was amplified as a control.

For human RBM10v1 and RBM10v2, the forward primer RBM10A, 5’-AGGGCAAGCATGACTATGA-3’ and reverse primer RBM10B, 5’-GTGGAGAGCTGGATGAAGG-3’ were used as described by Martinez-Arribas et.al., (30) For human RBM5 (LUCA-15), the forward primer RBM5, LU15 (2) 5’-GACTACCGAGACTATGACAGT-3’ and reverse RBM5, LU15(3) 5’-AGAGGACAGCTGCACAAATGC-3’ were used as described by Rintala-Maki et.al (31). For human exon 6-skipped Fas 5-7, the forward primer CCAAGTGCAAAGAGGAAGTGA and reverse primer GGAGATTCATGAGAACCTTGG were used, and for human exon 6-included Fas 5-6-7, the forward primer TGCAAAGAGGAAGGATCCAG and the reverse primer TTCTGTGTTTCTGCATGTTTT were used as described by (11). For human GAPDH, the forward primer 5’-AAGGCTGAGAACGGGAAGCTTGTCATCAAT-3’ and reverse primer 5’-TTCCCGTCTAGCTCAGGGATGACCTTGCCC-3’ were used as described by Nishio et.al., (32). The amplified cDNA products were separated on agarose gels (1.8 % w/v) containing ethidium bromide (0.1 µg/ml) and visualized by UV-exposure. The images of the amplified cDNA bands were captured by Olympus μ-7000 digital camera.

### Sequence aliment analysis of RBMs

The sequences from Homo sapiens RBM10v1 (NP_005667), RBM10v2 (NP_690595) and RBM5 (NP_005769) were aligned using the Cluster W multiple aliment algorithm (Vector NTI suite 7.0 AlignX). The carboxyl terminal 231 residues of RBM10v2, RBM10v1 and RBM5 were aligned and the highly conserved motifs were indicated.

## Results

### High expression levels of RBM10v1 and RBM5 in COS-7 and HeLa cells

COS-7, HeLa, A549 and HK-2 cells differentially expressed RBM10v1, RBM10v2 and RBM5 proteins as shown in Fig. 1A. HeLa and COS-7 cells expressed a higher level of RBM10v1and RBM5. A549 cells expressed a low level of RBM5 as previously described (33,34). HK-2 cells expressed the lowest levels of the RBMs. A549, HeLa and COS-7cells expressed RBM10v1 protein at twofold, fourfold and twelve-fold more than HK2 cells as shown in Fig.1A-a. A549, HeLa and COS-7 cells expressed a twofold, fourfold and six fold more amount of RBM10v1 over the RBM10v2 levels, respectively. A549, HeLa and COS-7 cells expressed a twofold, fourfold and sevenfold more RBM5, respectively, compared to HK-2 cells as shown in Fig. 1A-b. The nuclear extract of HeLa cells contained a twofold more RBM10v1 and a fourfold more RBM5 proteins compared to those of young TIG3S fetal fibroblasts (Fig. 1A-d). The overexpressions of the RBMs of the transformed cells are a novel finding but the molecular mechanism has not been clarified.

**Fig. 1.**
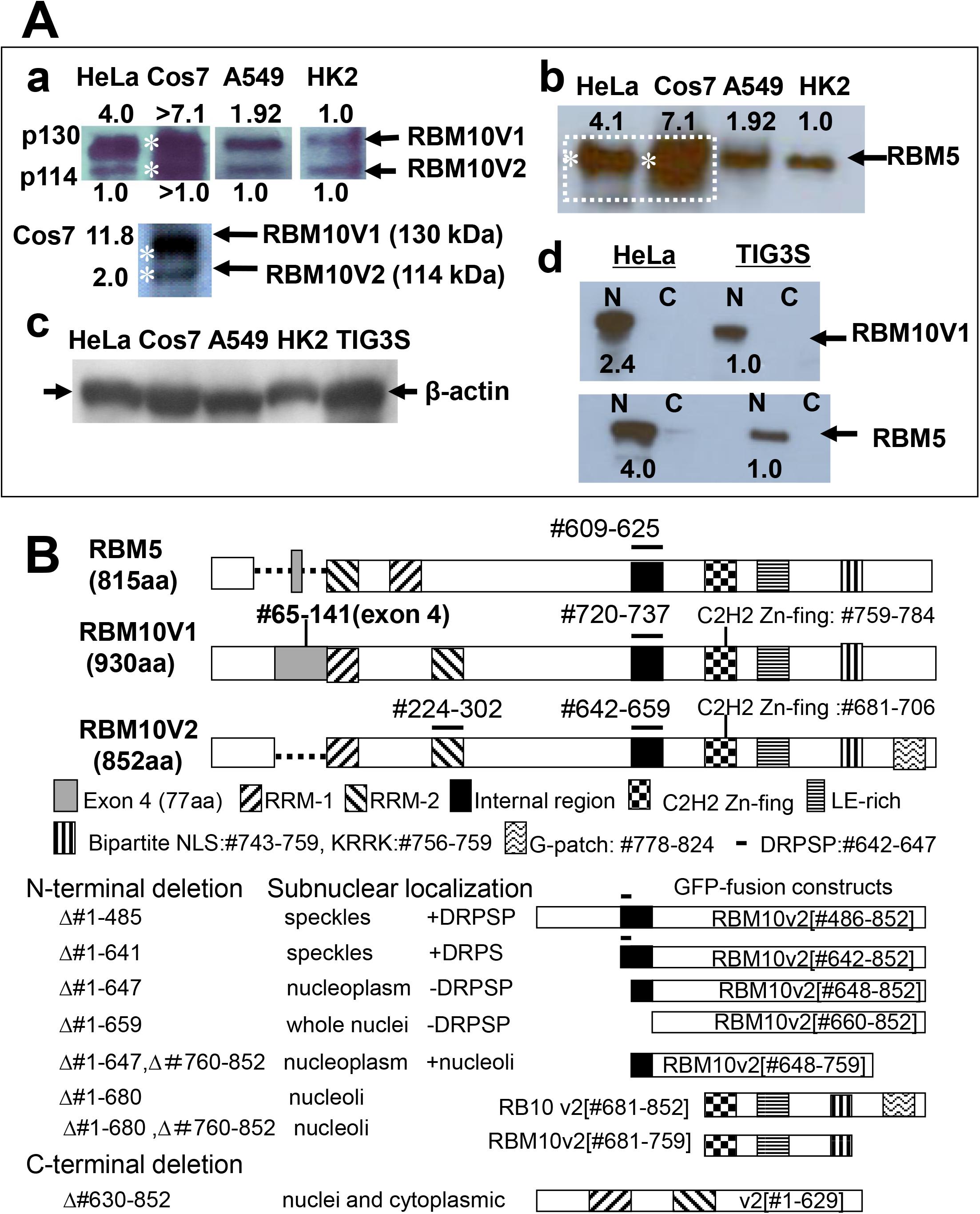
(A) RBM10 (v1and v2) and RBM5 protein levels in various cell lines. The whole cell lysates of COS-7, HeLa, A549, and HK2 cells (10μg of the protein) were separated on 8% SDS-PAGE. RBM10 and RBM5 protein were detected by Western blotting. The integrated optical densities of the specific protein bands were calculated by Image-Pro Plus software. The numbers above or below the bands indicate the relative protein levels of RBMs. RBM10v1 and RBM10v2 protein were detected in the lysates of HeLa, A549 and HK-2 cells (A-a). COS-7 cells expressed the higher amounts of RBM10 as indicated with white asterisks and thereby the RBM10 proteins were separately examined as shown in the lower panel. The RBM5 proteins were detected in the four cell lines (A-b). β-actin was compared as the loading control (A-c). The nuclear (N) and cytoplasmic (C) proteins (3 μg each) of HeLa and TIG3S (human fetal fibroblasts) cells were subjected to SDS-PAGE and Western blotting (A-d). (B) RRM1, RRM2, exon 4, and the highly conserved common motifs of three RBMs are illustrated. The constructs of the full length RBMs and the amino or carboxyl terminally deleted RBM10v2 are shown. DRPSPP: 6 amino acids residues of RBM10v2[#642-647] is critical for the nuclear speckles localization. Subnuclear localization of the each constructs is identified as follows: speckles, nucleoplasm, whole nucleus or nucleolus.

A positive correlation of the expression of both RBM10v1 and RBM5 was reported in human tumor tissues previously (35). In addition, strongly upregulated RBM mRNAs including RBM10 were found in nine cancers (36). Therefore, the higher protein levels of RBM10v1 and RBM5 seemed to be a characteristic of the highly proliferative tumor cells.

### Effects of antitumor agents on the stability of RBM proteins

Unstressed HeLa cells expressed the tiny RBM10 nuclear granules (25). Under normal culture condition, the nuclear speckles were not prominent in the actively growing A549 cells (Fig.2A). Hence, COS-7 cells were transfected with the expression vector of RBM10v1-GFP and N-terminally truncated RBM10v2 [#482-852]-RFP, and thereby the prominent nuclear speckles could be observed as shown in Fig. 2B. This result suggests that RBM10v2 [#482-852] preserves the structural elements for the nuclear speckles localization as discussed later.

**Fig. 2.**
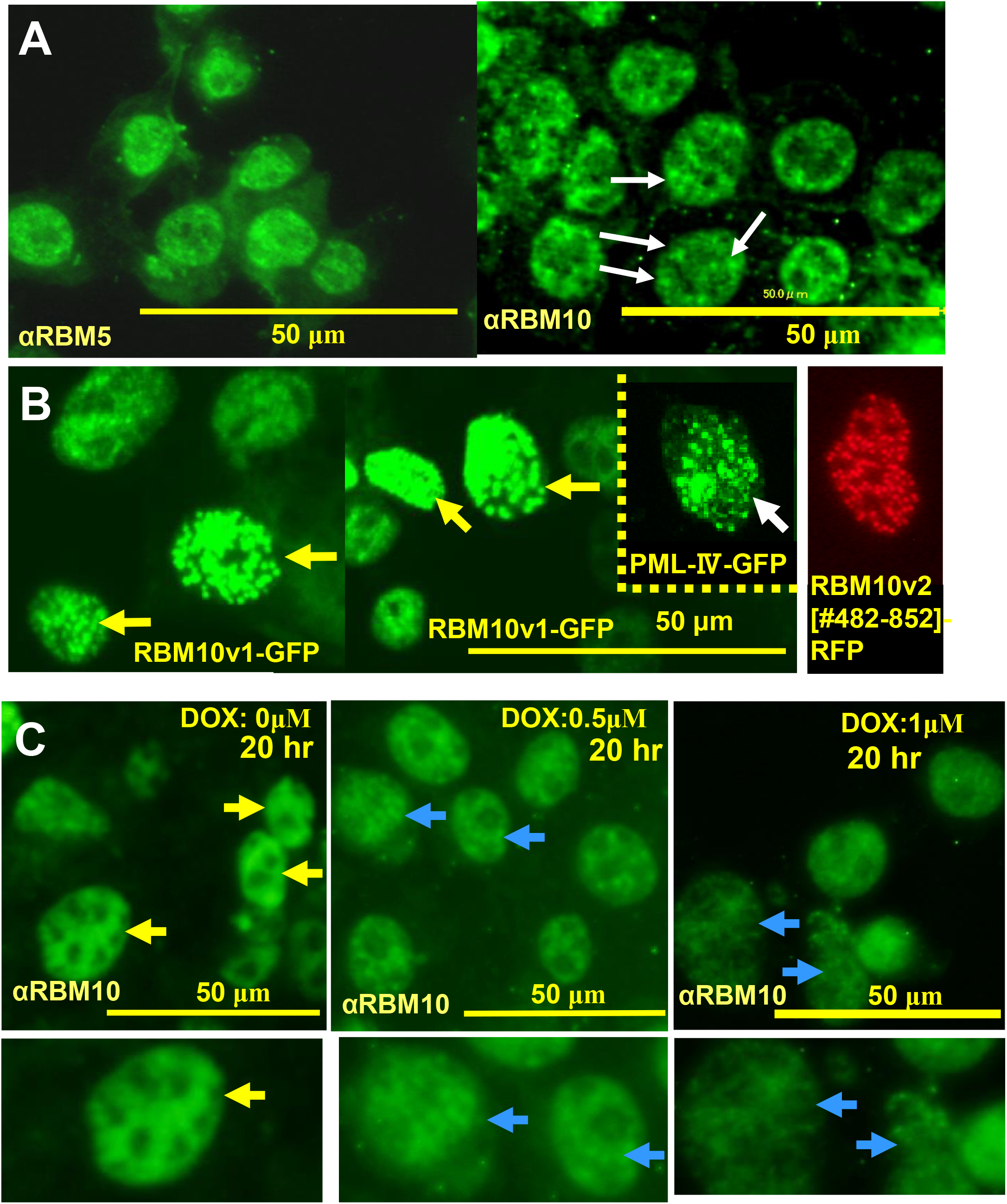
(A) Nuclear localization of the endogenous RBM5 and RBM10 in human lung carcinoma A549 cells. A549 cells were stained with the rabbit anti-RBM10 or anti-RBM5 serum and the secondary Alexa ^488^ labeled anti-rabbit IgG. The nuclear speckles of RBM10 are indicated by white arrows. (B) Nuclear speckles of RBM10v1-GFP are indicated by thick arrows and nuclear bodies of PML-Ⅳ-GFP are indicated by a thick arrow in the insert. The nuclear speckles of the N-terminally truncated RBM10v2[#482-852]-RFP are shown in the right panel. (C) A549 cells were exposed to 0.3 µM or 1 µM DOX for 20 hours and stained with the primary rabbit anti-RBM10 antibody and the secondary Alexa ^488^-labeled anti-rabbit IgG. The signal intensities of the nuclear RBM10 were examined by fluorescence microscopy as indicated by yellow (control) or blue arrows (DOX).

The effects of DOX and AcD on the expression of RBM10 and RBM5 were characterized. At first, A549 cells were exposed to DOX at a low therapeutic dose for 20 hours, and then the RBM10 immunofluorescent signals were examined. The treatment with 0.5 µM or 1 µM DOX decreased the RBM10 immunofluorescent signals as indicated by blue arrows in the middle and right panels of Fig.2C. In contrast, the non-treated cells kept the higher RBM10 signals as shown in the left panels of Fig. 2C. Interestingly, the DOX exposure did not promote the RBM10 nuclear speckles in A549 cells.

The drug sensitivity of the RBMs in COS-7 cells was quantitatively investigated as follows. The RBM proteins were examined after exposure to 1 µM DOX or 1 µM AcD as shown in Fig.3A-a and supplement Fig.S1A. Upon a 6 hour exposure to DOX, the RBM10v1 protein significantly decreased to 30% of the control level. RBM5 and β-actin proteins maintained at 96% and 85% respectively, of the control levels. Upon a 16 hour exposure to DOX, RBM10v1 protein was impaired to 3% of the control level (Fig.3A-a), but RBM10 mRNA remained at 66 % of the control level (Fig.3B). RBM5 and β-actin proteins remained at 68 and 47 % respectively, of the control levels. Thus, DOX more potently impaired the stability of RBM10v1 protein. AcD potently impaired the stability of RBM5 as follows: upon a 6 hour exposure to AcD, RBM10v1 protein remained at 53% of the control level, in contrast, RBM5 protein decreased to 3% of the control level as shown in Fig.3A-a. Upon a 16 hour exposure to AcD, RBM10v1 and RBM10v2 mRNAs were mostly impaired as shown in the upper panel of Fig.3B, whereas RBM5 mRNA considerably maintained at 34% of the control level.

**Fig. 3.**
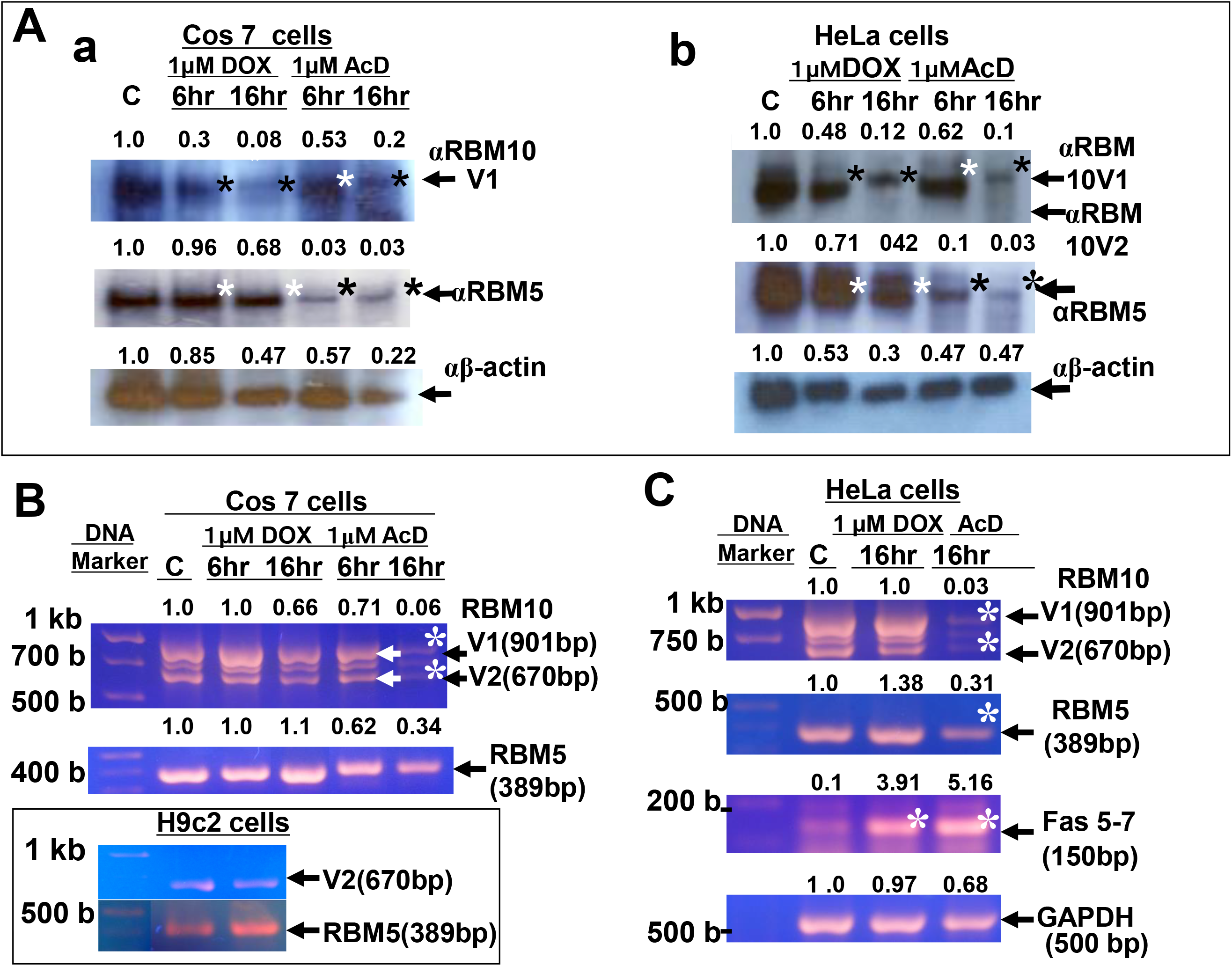
RBM10 and RBM5 protein levels of COS-7 or HeLa cells exposed to DOX or AcD. (A) COS-7 (left panel a) and HeLa cells (right panel b) were exposed to 1 µM DOX or 1 µM AcD for 6 hours and 16 hours. The lysates (10 μg) were subjected to Western blotting and the protein levels of RBM10, RBM5 and β-actin were examined. Relative band intensities for the specific protein bands were indicated (A) and those 3D bar plots were shown in supplement Fig.S1. Black asterisks show the significant reduction of RBM10 or RBM5 protein. White asterisks show the significant resistance of the RBM proteins. (B) RBM10 mRNAs of COS-7 cells exposed to DOX or AcD for 6 and 16 hours. Non-treated cells (indicated as C) or H9c2 cells were cultured without drugs. (C) RBM10 and Fas mRNAs of HeLa cells exposed to DOX or AcD for 16 hours (right panel). The total RNA was reverse transcribed and subjected to RTPCR. The endogenous mRNA levels of RBM10v1 (901bp), RBM10v2 (670bp), RBM5 (389bp), Fas 5-7 (150bp) or GAPDH (500bp) were determined by Image-Pro Plus software. The middle band between the both RBM10 variants, RBM10v3 was rarely observed. The numbers above the cDNA bands show the relative mRNA levels. Significant reduction or increase was indicated by white asterisk.

Next, HeLa cells were examined as follows. Upon a 6 and 16 hour exposure to DOX, RBM10v1 protein decreased to 48% and 12% of the control level, respectively as shown in Fig.3A-b and supplement Fig.S1B. Upon a 6 and16 hour exposure to AcD, RBM5 protein decreased to 10% and 3% of the control level, respectively as shown in Fig.3A-b. Hence, RBM5 protein was several folds more sensitive to AcD in contrast to RBM10v1. Upon a 16 hour exposure to AcD, RBM5 and RBM10 proteins were impaired completely, although the exon 6-skipped Fas mRNA (Fas 5-7) was mostly expressed (Fig.3C). PTB (polypyrimidine tract binding protein), HuR (anti-apoptotic regulator) and the N-terminal fragment of U2AF65 might simultaneously involve in the skipping of Fas exon 6 in the apoptosis-induced cells as reported previously (37–39). COS-7 and HeLa cells expressed RBM10 (RBM10v1>RBM10v2) and RBM5 mRNAs. Interestingly, H9c2 cells expressed the RBM10v2 and RBM5 mRNAs as shown in Fig.3B.

As noted above, DOX and AcD differentially affected the stability of RBM10 and RBM5. It is known that DOX and AcD activate caspase-2 and caspase-3 in normal or tumor cells. Thereby the RBMs become sensitive to proteolysis in circumstances of cancer chemotherapy. Accordingly, the putative caspase cleavage sites of the RBMs were searched by Expasy PeptideCutter tool that predicts protease sites (40). The results showed no caspase cleavage site of RBM10 and one caspase-1 cleavage site of RBM5 at #90D (YRH**D**ISDE), which suggests its higher sensitivity to AcD.

Proteasome-dependent nuclear protein degradation occurs in the distinct nuclear domains that overlap with ubiquitin, splicing speckles or PML bodies (41). Active PA28γ-proteasome complexes localize in the nuclear speckles and control the nuclear traffiking of splicing factors though selective proteolysis(42). DOX-mediated activation of ubiquitin proteasome system (UPS) was surveyed in cardiomyocytes or K562 erythroleukemic cells (18,43,44) and in vivo or in vitro (45). DOX induced the DNA damage of H1299 non-small cell lung carcinoma cells and triggered the increase of proteasome activity, of which caspase-like and chymotrypsin-like activities were 30% higher than non-treated cells (46).

Small ubiquitin-like modifier (SUMO) is identified as a targeting signal for ubiquitylation and ubiquitin-dependent degradation as reviewed by Miteva et al., (47). Interestingly, DOX specifically and significantly accelerated the proteasome-mediated proteolysis of transcription coactivator p300 and transcription factor NFAT5 in cardiomyocytes (48,49). Hence, I applied the SUMO prediction algorithm (50,51) to predict the sumoylation sites of RBM10v2, RBM5, NFAT5 and p300. The algorithm revealed that RBM10v2 (or RBM10v1) has 3 sumoylation sites including lysine residues at #54K, #163K and #458K and three SUMO-interaction motifs (SIMs) at #254-258, #681-685 and #806-810 as described in Table1. In contrast, RBM5 has one sumoylation site at #181Kand two SIMs at #262-266 and #647-651. Interestingly, p300 or NFAT5 has six or seven sumoylation sites, respectively and two SIMs. Taken together, the bioinformatics analysis supports that proteasome mediates the degradation of RBM10, NFAT5 and p300.

### Actinomycin D promotes the targeting of RBM10v2 and RBM5 to the nuclear speckles

COS-7 cells were transfected with the pRBM10v2-RFP construct, cultured for 48 hours and followed in the medium containing DOX. Red fluorescence of the RBM10v2-RFP and green fluorescence imaging of the endogenous RBM10 were examined. RBM10v2-RFP diffusely localized in the nucleoplasm of many cells (Fig. 4A, top panel). Upon an 8 hour exposure to 0.5 µM or 1 µM DOX, a very few of the blurred nuclear speckles of RBM10v2-RFP were evidenced as marked with white arrows (Fig. 4B or Fig. 4C). The result indicates that DOX induced the proteolysis of the RBM10v2 protein.

**Fig. 4.**
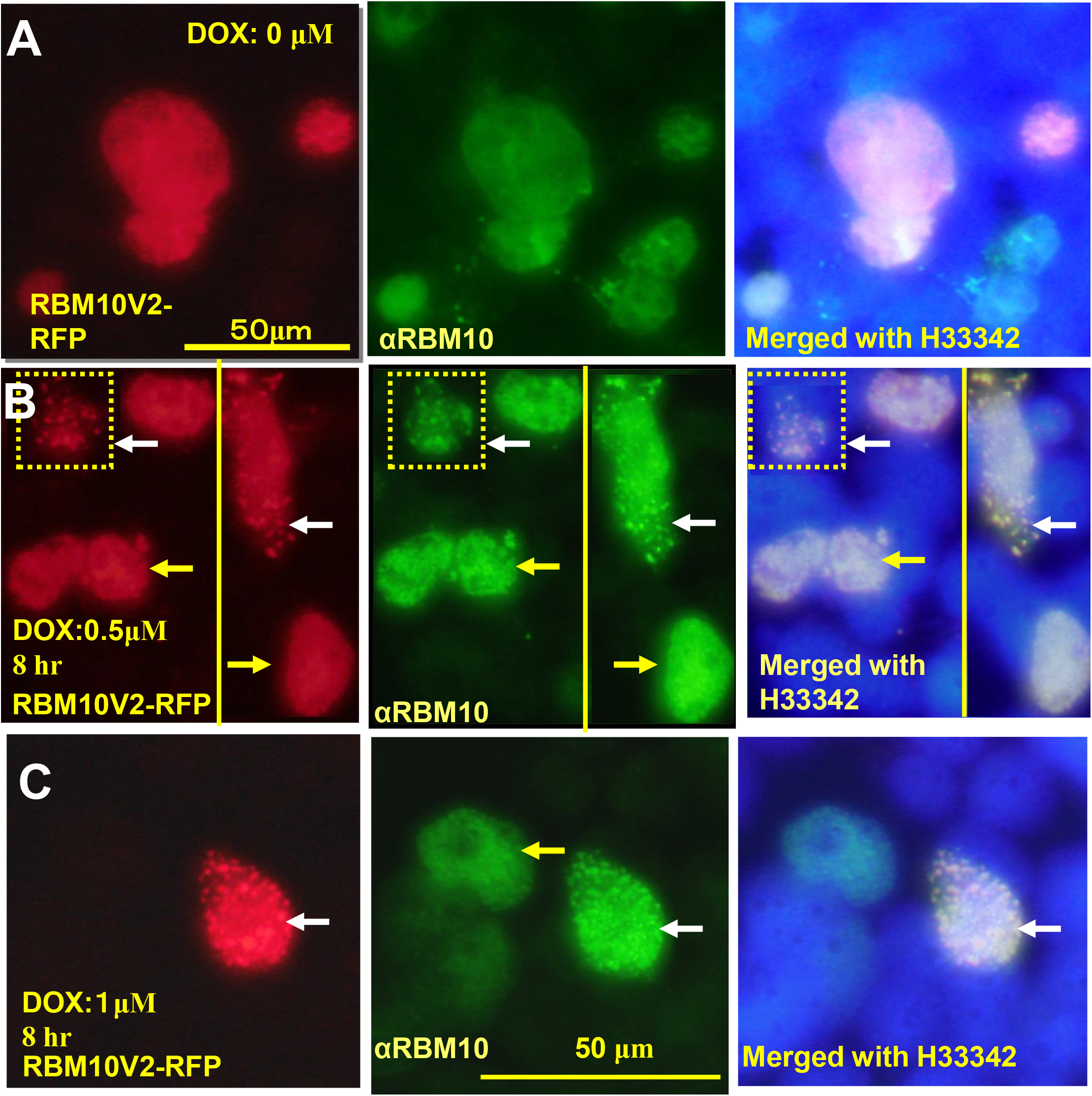
Nucleoplasmic and nuclear speckles localization of RBM10. (A) Nucleoplasmic localization of RBM10v2-RFP and endogenous RBM10 in COS-7 cells (top panel). RBM10 were detected with the rabbit anti-RBM10 and secondary Alexa^488^-labeled anti-rabbit IgG. COS-7 cells grown in a 4-well culture chamber slide were transfected with 1 μg of pRBM10v2-RFP with 5 μl of lipofectoamin and then cultured for 48 hours. The endogenous RBM10 and RBM10v2-RFP was examined after 8 hours-exposure of 0.5 μM DOX (B) or 1 µM DOX (C). The nuclei were labeled with H33342. The green and red fluorescence images of the RBMs were captured with fluorescence microscopy.

COS-7 cells were transfected with the expression construct of pRBM10v2-RFP, of which tiny nuclear speckles were evidenced as shown in Fig.5A. The nuclear speckles of RBM10v2-RFP colocalized with the green fluorescence imaging with the rabbit anti-RBM5 and anti-rabbit IgG-Alexa^488^. Upon a 4 hour exposure of the cells to 1 µM AcD, the prominent nuclear speckles of RBM10v2-GFP were promoted as indicated by yellow arrows in Fig. 5B. The nuclear speckles of RBM10v2-GFP remained after 20 hour-exposure to AcD (Fig.5B, lower panels). Furthermore, the endogenous RBM5 were detected with rabbit α-RBM5 antibody and Alexa^555^-laveled secondary anti-rabbit IgG. The red fluorescence imaging of the endogenous RBM5 successfully colocalized with the nuclear speckles of RBM10v2-GFP. The perfect colocalization of the RBM10v2-GFP (or RFP) and the endogenous RBM5 suggests their presence in the same nuclear speckles. In addition, A549 cells were transfected with the pRBM5-GFP construct, and RBM5-GFP localized diffusely in the nuclei (Fig.5C). Upon a 3 hour exposure to AcD, the targeting of RBM5-GFP to the nuclear speckles was promoted (Fig.5D).

**Fig. 5.**
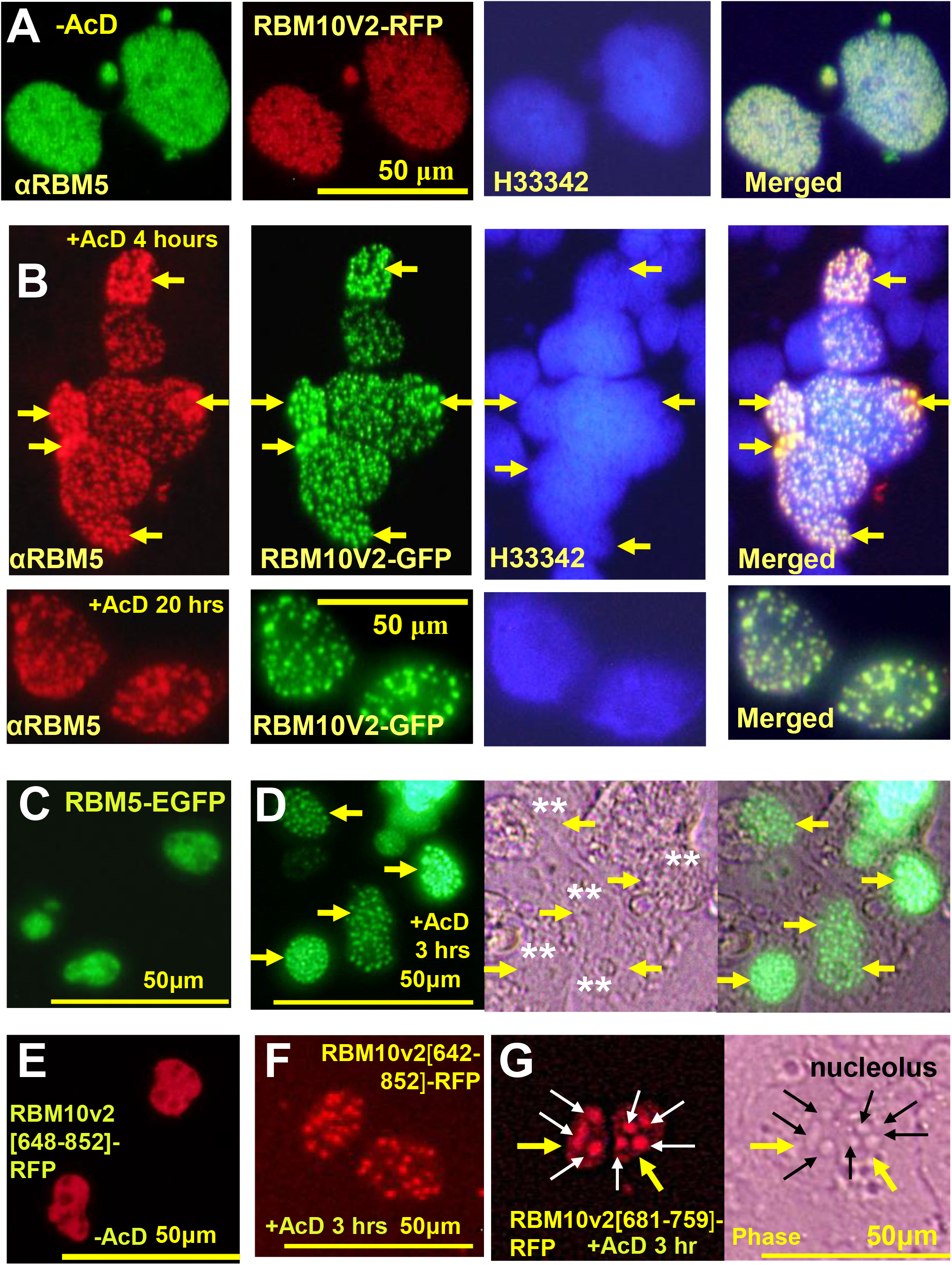
Nuclear speckles of endogenous RBM5 and exogenous RBM10v2-GFP (or RFP). COS-7 cells were transfected with pRBM10v2-RFP or pRBM10v2-GFP as described previously (top panel). The endogenous RBM5 were detected with the anti-RBM5 and the secondary antibody Alexa^488^ or Alexa^555^ as described previously. The numerous small nuclear speckles of the endogenous RBM5 and RBM10v2-RFP could be expressed in COS-7 cells (A). Upon exposure to 1μM AcD for 4 or 20 hours (lower panels), the prominent nuclear speckles appeared as indicated as yellow arrows (B). The full length RBM5-GFP diffusely localized in the nucleoplasm of A549 cells by live cell microscopy (C). Addition of AcD for 3 hours promoted the targeting of RBM5-GFP to the nuclear speckles (yellow arrows) and double asterisks indicated the nuclei (D). The N-terminally truncated RBM10v2[#648-852]-RFP localized in the nucleoplasm of COS-7 cells by live microscopy (E). In A549 cells, the nuclear speckle localization of RBM10v2[#642-852]-RFP was not affected by AcD (F). The nucleolar targeting of the truncated RBM10v2[#681-759]-RFP and the nucleoli of A549 cells were not affected by AcD (G). The nucleoli were indicated by thin black arrows (phase contrast image).

Structural elements of RBM10 targeting to the nuclear speckles were explored as described below. The expression vectors of GFP-fusion of RBM10v2 comprising the N-terminal [#1-629] and RFP-fusion of RBM10v2 comprising the C-terminal [#482-852],[#642-852], [#648-852],[#660-852],[#648-759], [#681-852] and [#681-759] were constructed as shown in Fig.1B. The expression vectors were expressed in COS-7 cells unless otherwise noted. At first, RBM10v2 [#482-852]-RFP localized in the nuclear speckles as previously noted (Fig.2B). Similarly, RBM10v2 [#642-852]-RFP including the C-terminal 213 amino acids, was targeted to the nuclear speckles (Fig.5F). Upon the deletion of N-terminal 6 amino acids (*DR****P****SPP*),the resulting RBM10v2[#648-852]-RFP mostly localized in the nucleoplasm (Fig.5E). Consequently, *DR****P****SPP*[#642-647] is critical for the speckle localization in COS-7 cell system. After 3 hour exposure to AcD, the nuclear speckles of RBM10v2 [#642-852]-RFP remained unchanged as similar as none-treated (Fig.5F). Whereas, RBM10v2[#660-852]-RFP uniformly localized in the whole nuclei but RBM10v2 [#1-629]-GFP diffusely localized in the cytoplasm and nuclei (supplement Fig.S2-A and S2-C respectively). The above results suggest that RBM10v2 [#1-629] and RBM10v2 [#660-852] lost the ability of targeting to nuclear speckles.

The carboxyl terminal 231 amino acids of RBM10v2, RBM10v1 and RBM5 include the highly conserved motifs, as shown in supplement Fig.S4. The C-terminal region include a SIM <u>**LACLL**</u> [#681-685] at C2H2 Zinc-finger [#681-706], a LE-rich (leucine and glutamic acid rich) [#710-727], a region [#734-759] containing bipartite NLS [#743-759] and G-patch motif (glycine-rich RNA binding domain) [#778-824]. Primarily, the region [#642**-**659] (*DR<u>**P**</u>SPP<u>**RGLVAAYSG**E**SD**</u>*) significantly affects the nuclear speckles targeting of RBM10. Importantly, human RBM5 has the quite similar sequence at [#609-625] (*EN<u>**P**</u>*-LK<u>***RGLVAAYSG**D**SD***</u>). The completely or positively identical amino acid residues are *underlined bold italic* or *italic*, respectively. Thereby RBM10 and RBM5 conserve the region that plays a potent role in the targeting to the nuclear speckles.

Interestingly, RBM10v2-[#681-852]-RFP mostly localized in the nucleoli (supplement Fig.S2-B). RBM10v2 [#681-759]-RFP localized in the nucleoli and remained unchanged upon a 3 hour exposure to AcD as shown in Fig.5G. Notably the nucleolar targeting of both RBM10v2 [#681-852] and RBM10v2 [#681-759] was quite different from the nucleoplasm targeting of RBM10v2 [#648-852] and nuclear speckles targeting of RBM10v2 [#642-852]. Hence, it is likely that RBM10v2[#681-759, #681-852] has a N-terminally exposed SIM **LACLL** and a nucleolar targeting element, whereas the N-terminal region RBM10v2[#648-680] seems to mask the nucleolar targeting element.

The motif of nucleolar localization signal contains (R/K) (R/K) X (RK) or (R/K) X(R/K) (R/K) as reported previously (52). Histone H2B has a nucleolar retention signal sequence, KKR**KR**S**RK** (53). Moreover sumoylation of nucleoplasmin/B23 is necessary for its nucleolar residency and function (54). RBM10v2 has a bipartite NLS motif [#743-759] **RR**E**K**YGIPEPPEP**KRRK**. As described above, the basic amino acid cluster **KRRK** seems to be a motif of nucleolar retention signal of RBM10v2 [#681-759, #681-852]-RFP. In addition, RBM10v2 [#681-852] and RBM10v2 [#681-759] have a SIM **LACLL** [#681-685]. Thereby, the **KRRK** signal and SIM of RBM10v2 [#681-759] may interact with the sumoylated nucleolar components.

## Discussion

### Expression of RBM10 and RBM5 in human tumor cells

RBM10v1 and RBM10v2 were detected among the several human tumor cell lines. COS-7 and HeLa cells expressed RBM10v1 and RBM5 at higher levels. Notably, H9c2 rat cardiomyoblasts expressed mostly RBM10v2 mRNA. The result is consistent with the significant high expression of RBM10v2 protein in H9c2 cells (55). Taken together, it is likely that the RBM10v1 and RBM10v2 are differentially expressed in COS-7, HeLa and H9c2 cells, respectively as recently reported (56).

### Effects of antitumor agents to RBM10 and RBM5

DOX enhanced the proteasome or UPS-mediated degradation of cardiac troponins, cardiac transcription factors and survival factors (18,49,57). DOX specifically enhanced the degradation of nuclear protein p300 and nuclear factor-activated T cell 5 in the cultured cardiomyocytes (48,49). Similarly, the present study revealed that RBM10 of the DOX-exposed COS-7 cells was highly sensitive to proteolysis.

In HeLa or COS-7 cells, a 3 hour exposure to AcD induces the RBM6 nuclear speckles (24). Upon a 24 hour exposure to 0.8 µM AcD, HeLa cells decreased the endogenous snRNP and SC35 proteins (58). In the present investigation, COS-7 cells were transfected with the expression vector of RBM10-RFP and followed with a short time-exposure to AcD, resulting in the prominent nuclear speckles of the endogenous RBM5 and RBM10v2-RFP.

Mitoxantrone (MTX) is a structurally DOX-related topoisomerase II inhibitor. When A549 cells were treated with 2.5 µM MTX for 24 hours, the RBM10v1 and RBM10v2 mRNA mostly disappeared, but RBM5 mRNA mostly remained unchanged (data not shown). MTX also differentially triggered the decay of RBM10 mRNA.

### Nuclear speckles of RBM10 and PML nuclear bodies

The nuclear speckles of RBM10 and RBM5 were more prominent in the AcD-exposed cells. Next, the structural elements of the RBM10 nuclear speckles were investigated. COS-7 cells were transfected with the N-terminally truncated RBM10v2-RFP construct that lacks the N-terminal 481 or 641 residues. The RFP-fusion of RBM10v2 [#482-852] or RBM10v2 [#642-852] localized in the prominent nuclear speckles under the normal culture conditions. In addition, A549 cells were transfected with the pRBM10v2 [#1-629]-NLS-GFP construct. The N-terminal 629 residues included RRMs, one SIM in the RRM2, three sumoylation sites and NLS (PKKKRKV) of SV40LG. Regardless of exposure to AcD, RBM10v2 [#1-629]-GFP diffusely localized in the cytoplasm, but neither localized in the nuclear speckles nor the nucleoli (supplement Fig.S2-C) even after AcD exposure. Hence, the N-terminal 629 residues do not include the targeting element of the nuclear speckles. In addition, a recent investigation reported that KEKE motif [#549-568] and Zing finger C_2_H_2_ [681-706] are the nuclear speckles targeting sequences in ARL cells (rat liver epithelial cells) (59). The differential localization of the truncated RBM10 might occur in the above two cell systems.

The conserved regions, RBM10v2 [#648-663]<u>***RGLVAAYSG**E**SD***</u>S<u>***EEE***</u> and [#672-680]***EEKL***T***DW***Q***K*** are unique and thereby their deletions resulted in the nucleolar translocation of the truncated RBM10v2 [#681-759]. The region RBM10v2 [#648-680] seems to have an inhibitory effect on the nucleolar targeting. Taken together, it is proposed that RBM10v2 [#642-680] has the structural elements that affect the translocation to the nuclear speckles, nucleoplasm or nucleolus.

Nuclear speckles are enriched with pre-mRNA splicing/processing factors and transcription repressors, but do not represent major sites of transcription and splicing. A long non coding regulatory RNA, MALAT1 involves in the cycling of the SR splicing factors between nuclear speckles and spliceosome and modulate alternative splicing (60). Wilms tumor suppressor WT1 (+KTS) interacts with the splicing factor U2AF65 (61) and undergoes nuclear speckles localization which requires the sumoylation of other proteins (62,63). PML and its sumoylation are essential for the formation of PML nuclear bodies as previously reviewed (47). Human cytomegalovirus (HCMV) immediate-early 1 (IE1) protein undergoes sumoylation which is required for the activity and efficient HCMV replication (64,65). IE1 interacts with sumoylated PML and prevent or remove SUMO-adducts to disrupt PML nuclear bodies (66). Interestingly, IE1 protein interacts with some dozens of nuclear proteins, and among them, seven are RNA binding proteins including RBM10 and RBM5 (67).

PML-Ⅵ and PML-Ⅳare 560 amino acids and 633 amino acids proteins, respectively, and both PML isoforms have multiple sumoylation sites (68). The carboxyl terminus of PML-Ⅳinteracts with the core domain of p53, of which nuclear body localization depends on the sumoylation of PML-Ⅳ(69). The RING motif of PML and the sumoylation of Sp1 are essential for the recruitment of Sp1 into PML nuclear bodies. The SIM of PML also plays an important role in this process (70). Thus, sumoylation and SIM should mediate the protein-protein interactions between PML, Sp1, p53 and various nuclear proteins including RBM10 and RBM5.

Of course, RBM10v2 shares the low sequence identities, 11.4% and 9.8% with PML-Ⅳ and PML-Ⅵ, respectively. The SUMO prediction algorithm predicted one sumoylation site (#386K) of p53 and a SIM (#143-147: **VQLWV)** at the p53 core domain, a specific fourth sumoylation site (#616K) of PML-Ⅳ and a SIM (#550-560: **VVVIS)** as described in Table 1. Interestingly, PML-Ⅵ lacks the above two motifs. Accordingly, I explored a certain interaction of RBM10 with PML-Ⅳ, and p53 as described below. In COS-7 cells, the coexpressed RBM10v2-RFP and PML-Ⅵ-GFP did not colocalize on the nuclear speckles or nuclear bodies (supplement Fig. S3-A). This suggests no specific interaction between RBM10v2 and PML-Ⅵ. However, RBM10v1-RFP and RBM5-GFP colocalized on the nuclear speckles (supplement Fig. S3-B), as previously described. Similarly, the nuclear speckles of RBM10v2-RFP and RBM10v2[#642-852]-RFP colocalized on the nuclear bodies of PML-Ⅳ-GFP (supplement Fig. S3-C). This demonstrated a certain interaction between the two proteins. Furthermore, p53-GFP colocalized with the nuclear speckles of RBM10v2-RFP in human fibroblasts (data not shown). Taken together, these results suggest that structurally unrelated proteins, RBM10, PML-Ⅳor p53 undergo sumoylation and interact through the attached SUMO and SIMs.

**Table 1.**
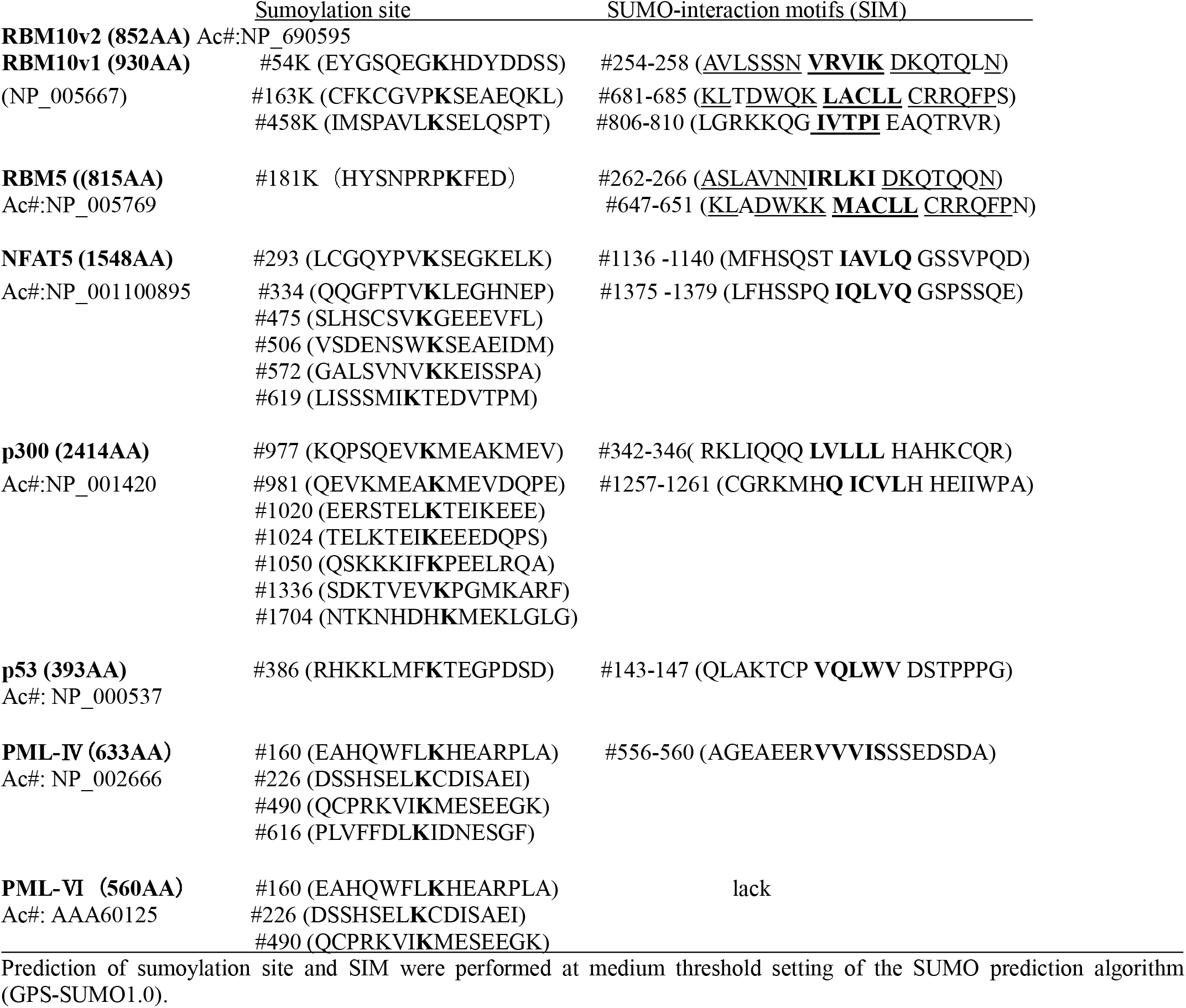
Sumoylation and SUMO-interaction sites in RBM10, RBM5, NFAT5 and p300.

## Ethical approval

This article does not contain any studies with animals performed by the author.

## Acknowledgements

This work was supported in Grant in Aid for Science Research (C) by the Japan Society for the Promotion of Science (17590159 to K.N).

## Abbreviations

AcD: Actinomycin D
DOX: Doxorubicin
GFP: green fluorescent protein
PML: promyelocytic leukemia protein
RBM: RNA binding motif (RBM) Protein
RBM5: RNA binding motif protein 5
RBM10: RNA binding motif protein 10
RRM: RNA recognition motif
RBM10v1: RBM10 variant 1
RBM10v2: RBM10 variant 2
VEGF: vascular endothelial cell growth factor
SUMO: small ubiquitin-like modifier
SIM: SUMO interaction motif

## Supplemental materials captions

**Supplement Fig. S1.** Differential sensitivities of RBM10 and RBM5 to Doxorubicin or AcD. COS-7 (A) or HeLa cells (B) were exposed to DOX or AcD for 6 and 16 hours as described previously. The protein levels of RBM10, RBM5 and β-actin were shown as graph bar plots. Black or red asterisks denote the resistance or sensitivity of the proteins, respectively. Differential drug sensitivities of RBM10 and RBM5 protein were demonstrated.

**Supplement Fig. S2** (A) Diffuse nuclear localization of RBM10v2 [#660-852]-RFP in COS-7 cells. (B) Nucleolar localization of RBM10v2[#681-852]-RFP in COS-7 cells. C) Diffuse cytoplasmic/nuclear localization of RBM10v2 [#1-629]-GFP in A549 cells. The cells were transfected with the expression vectors and observed by fluorescence-phase contrast live microscopy. The nuclear localization was indicated by yellow arrows. The nucleoli were indicated by white or black arrows (phase contrast image). RBM10v2 [#660-852]-RFP and RBM10v2[#1-629]-GFP localized neither on the nuclear speckles nor the nucleoli.

**Supplement Fig. S3** (A) Coexpression of RBM10v2-RFP and PML-Ⅵ-GFP in COS-7 cells by live microscopy. Nuclear speckles of RBM10v2 or nuclear bodies of PML-Ⅵ were indicated by yellows or blue arrows, respectively. No colocalization of RBM10v2-RFP and PML-Ⅵ-GFP was shown in a right panel. (B) Coexpression of RBM10v1-RFP and RBM5-GFP in COS-7 cells. The both nuclear speckles of RBM10v1 and RBM5 mostly colocalized as indicated by yellow arrows. (C) Coexpression of RBM10v2-RFP and PML-Ⅳ-GFP in COS-7 cells. The nuclear bodies of PML-Ⅳ-GFP (white arrows) partially colocalized with the nuclear speckles of RBM10v2-RFP (upper panels) or those of RBM10v2 [#642-852] (lower panels) indicated by yellow arrows.

**Supplement Fig.S4 Alignment** of carboxyl terminal 231 amino acids of human RBM10v2, RBM10v1 or RBM5. RBM10v2 [#622-852] and RBM10v1 [#700-930] have a 100% positive identities with RBM5 [#589-815]. Residues in the alignment are colored according to the following scheme. Black: non-similar residues, blue: consensus residue derived from a block of similar residues, red: consensus residue derived from a completely conserved residue, and green: residue weakly similar to consensus residue at given position. The highly conserved RBM10v2 [#642-659] are highlighted in yellow. DR**P**SP*P* [#642-647] is critical for the nuclear speckles. LACLL and IVTPI are SIM. KRRK is a nucleolar retention signal. The Zing finger [#681-706], LE (leucine and glutamic acid)-rich [#710-727], NLS [# 743-759] and G-patch motifs [#778-824] are highlighted in light blue color. Human RBM10v2 and rat RBM10v2 share 97.8% of positive identity and their C-terminal 252 residues are completely conserved.

