## Supplementary figures and images for "Differential effects of Doxorubicin and Actinomycin D on the stability of RNA binding proteins, RBM10 and RBM5: Actinomycin D promotes the nuclear speckles targeting of RBM10 and RBM5 through the novel structural elements"

### Supplemen Fig S3

supplement Fig. S3

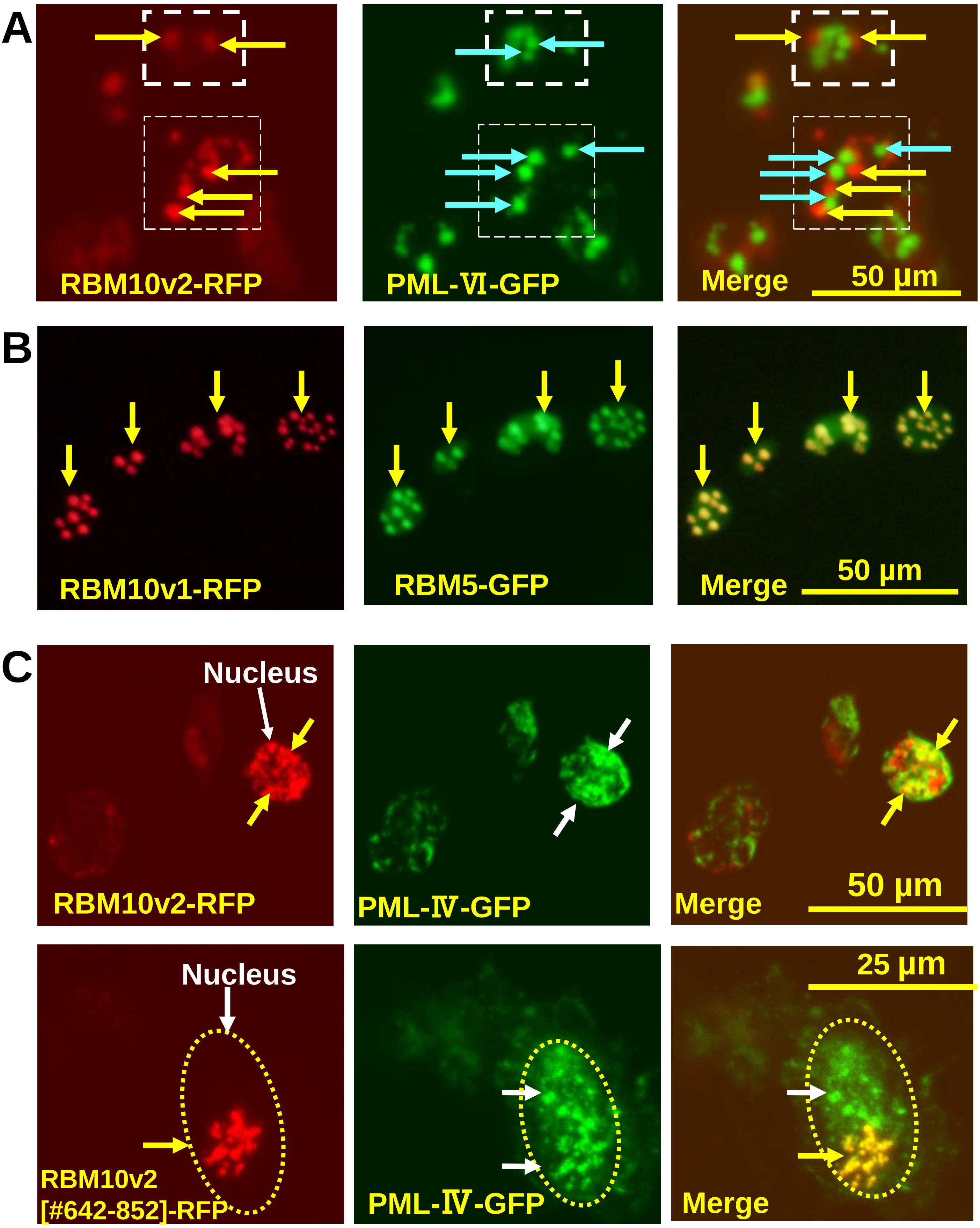

### Supplement Fig S1

Supplement Fig.S1

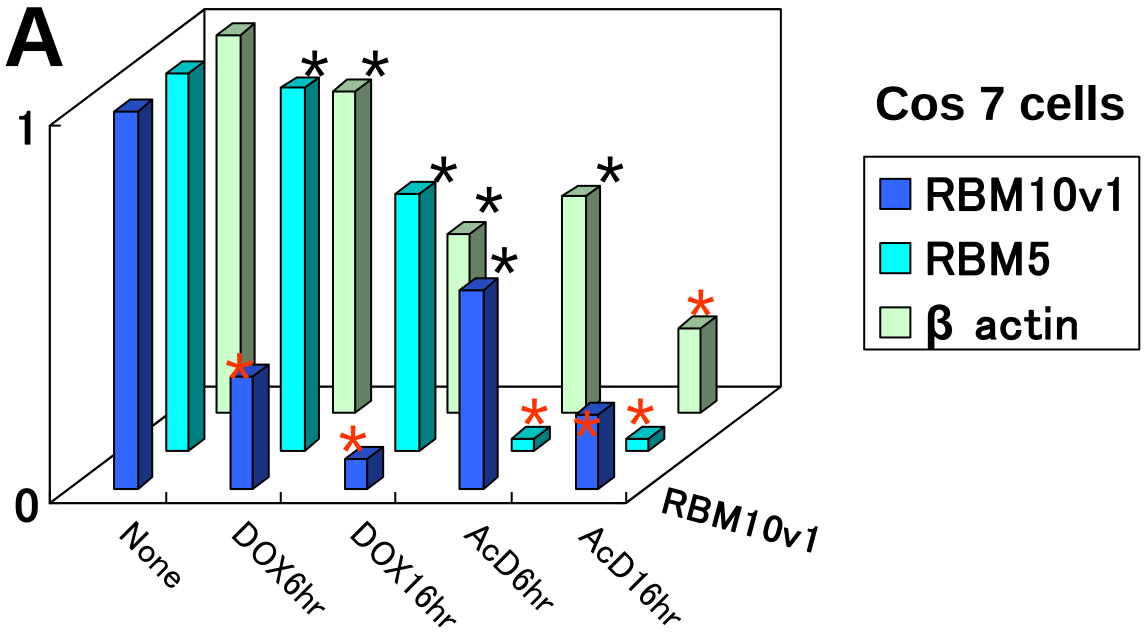


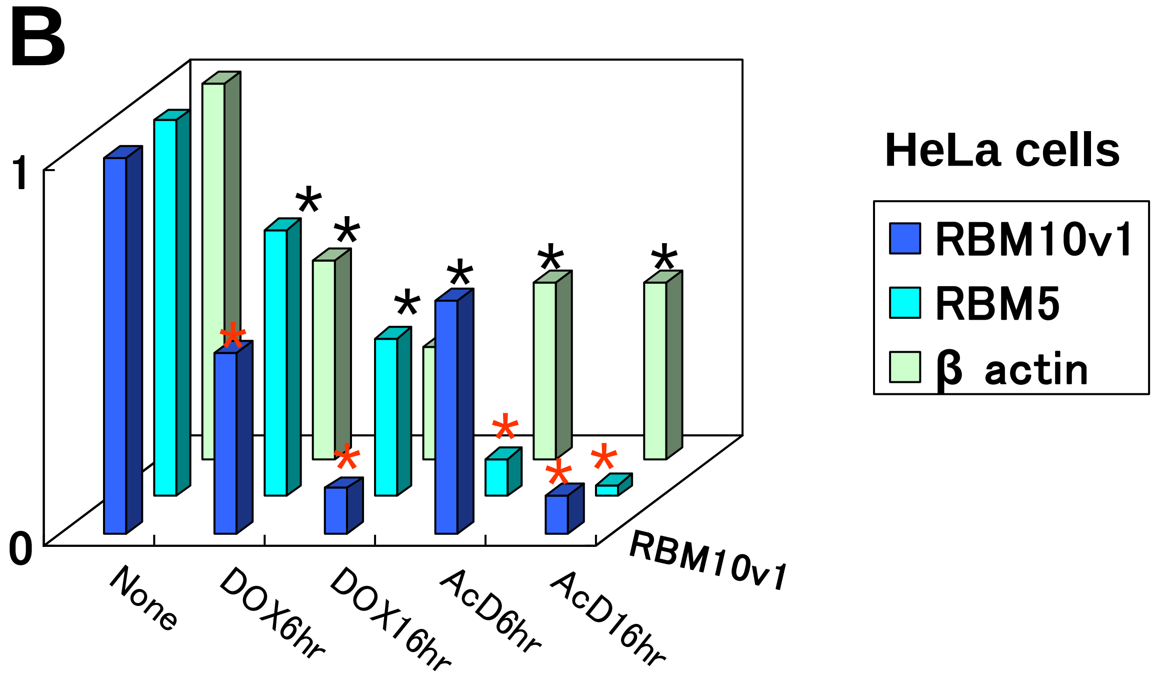

### Supplement Fig S2

Supplement Fig.S2

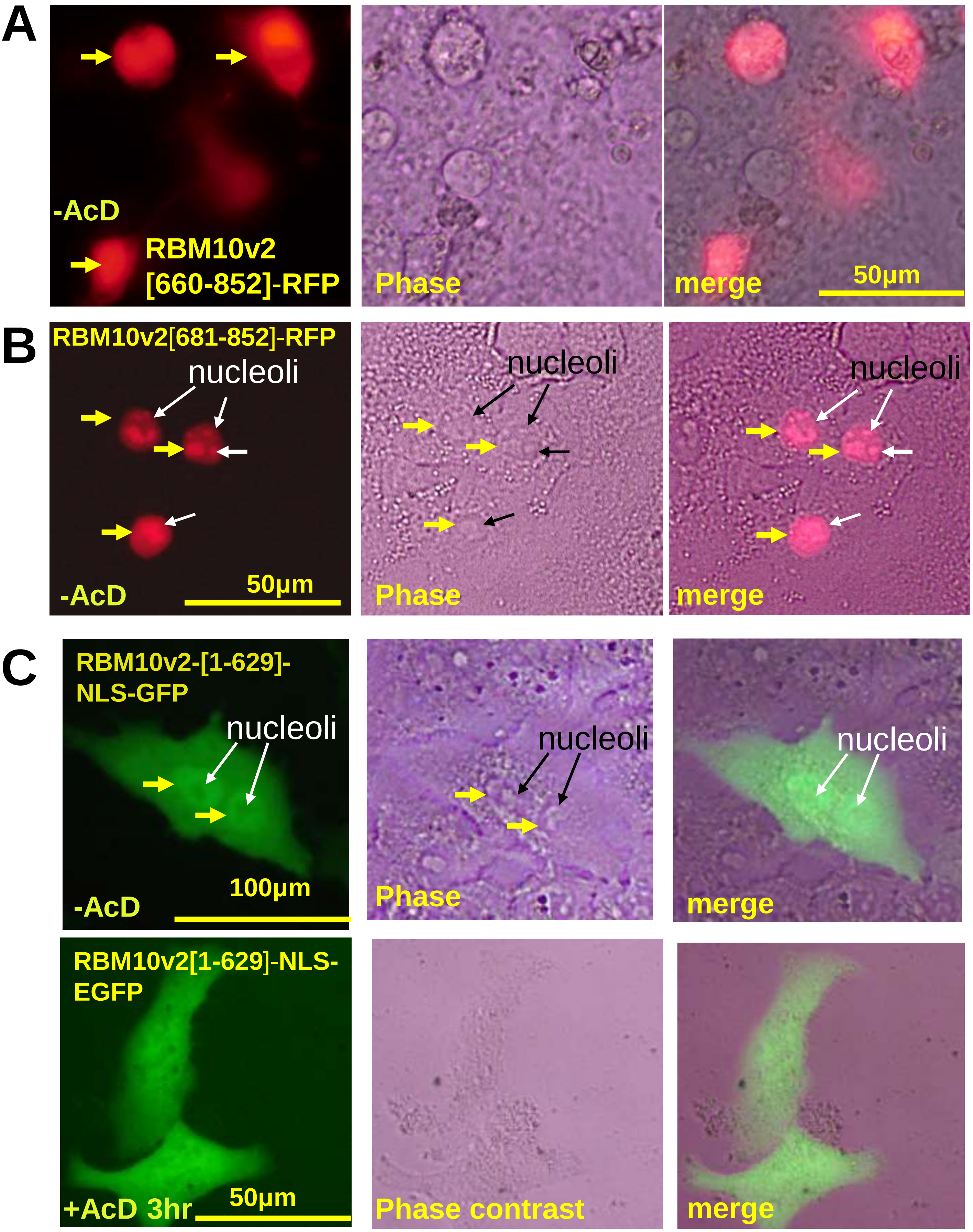
