## Supplement Fig s4 for "Differential effects of Doxorubicin and Actinomycin D on the stability of RNA binding proteins, RBM10 and RBM5: Actinomycin D promotes the nuclear speckles targeting of RBM10 and RBM5 through the novel structural elements"

```
                    622          |630          |640        |648              |660           |670
human RBM10v2 (622) KGALAERQHTSMDLPKLASDDRPSPPRGLVAAYSGESDSEEEQERGGPERE
human RBM10v1 (700) KGALAERQHTSMDLPKLASDDRPSPPRGLVAAYSGESDSEEEQERGGPERE
human RBM5    (589) KGALAERQQLIPELVRNGDEENP-LKRGLVAAYSGDSDNEEELVERLESEE

                    673        |680           |690             |700          |710            |720
human RBM10v2 (673) EKLTDWQKLACLLCRRQFPSKEALIRHQQLSGLHKQNLEIHRRAHLSENEL
human RBM10v1 (751) EKLTDWQKLACLLCRRQFPSKEALIRHQQLSGLHKQNLEIHRRAHLSENEL
human RBM5    (639) EKLADWKKMACLLCRRQFPNKDALVRHQQLSDLHKQNMDIYRRSRLSEQEL

                    724        |730           |740           |750          |759              |770
human RBM10v2 (724) EALEKNDMEQMKYRDRAAERREKYGIPEPPEPKRRKYGGISTASVDFEQPT
human RBM10v1 (802) EALEKNDMEQMKYRDRAAERREKYGIPEPPEPKRRKYGGISTASVDFEQPT
human RBM5    (690) EALELRERE-MKYRDRAAERREKYGIPEPPEPKRKKQ--FDAGTVNYEQPT

                    775     |780           |790             |800           |810             |820
human RBM10v2 (775) RDGLGSDNIGSRMLQAMGWKEGSGLGRKKQGIVTPIEAQTRVRGSGLGARG
human RBM10v1 (853) RDGLGSDNIGSRMLQAMGWKEGSGLGRKKQGIVTPIEAQTRVRGSGLGARG
human RBM5    (738) KDGIDHSNIGNKMLQAMGWREGSGLGRKCQGITAPIEAQVRLKGAGLGAKG

                    826   |830             |930              |852
human RBM10v2 (826) SSYGVTSTESYKETLHKTMVTRFNEAQ
human RBM10v1 (904) SSYGVTSTESYKETLHKTMVTRFNEAQ
human RBM5    (789) SAYGLSGADSYKDAVRKAMFARFTEME
```
